# Marsilea: An intuitive generalized visualization paradigm for complex datasets

**DOI:** 10.1101/2024.02.14.580236

**Authors:** Yimin Zheng, Zhihang Zheng, André F. Rendeiro, Edwin Cheung

## Abstract

Contemporary data visualization is challenged by the growing complexity and size of datasets, often comprising numerous interrelated features. Traditional visualization methods struggle to capture these complex relationships fully or are specialized to a domain requiring familiarity with multiple visualization tools. We introduce a novel and intuitive general visualization paradigm, termed “cross-layout visualization”, which integrates multiple plot types in a cross-like structure. This paradigm allows for a central main plot surrounded by secondary plots, each capable of layering additional features for enhanced context and understanding. To operationalize this paradigm, we present “Marsilea”, a Python library designed for creating complex visualizations with ease. Marsilea is notable for its modularity, diverse plot types, compatibility with various data formats, and is available in a coding-free web-based interface for users of all experience levels. We showcase its versatility and broad applicability by re-creating existing visualizations and creating novel visualizations that include elements such as heatmaps, sequence motifs, and set intersections that are typically beyond the scope of existing general visualization tools. The cross-layout paradigm, exemplified by Marsilea, offers a flexible, customizable, and intuitive approach to complex data visualization, promising to enhance data analysis across scientific domains.

## Main

Data visualization for research and data science faces significant current challenges, primarily due to the exponential growth and complexity of datasets. These datasets often include numerous features, making it difficult to display and interpret the information effectively. Traditional visualization techniques are frequently inadequate for these complex, multi-featured datasets, as they struggle to capture the intricate relationships and the full scope of the data. To mitigate these limitations, researchers have employed a variety of strategies. Dimensional reduction techniques such as Principal component analysis (PCA) or Uniform Manifold Approximation and Projection (UMAP)^1^ offer some relief but are limited in representing individual features or statistical details. Hence, researchers use multiple plots such as violin plots, bar plots, or heatmaps, each highlighting different aspects of the dataset. However, this approach fails to convey interrelations between features intuitively, leaving readers to examine multiple plots and captions back-and-forth to capture such interrelations themselves. Another strategy involves the creation of more complex visualizations, such as the complex heatmap^2^ for genomic data, multiple sequence alignment (MSA) plot for DNA/protein sequence comparison, upset plot^4^ for set intersection, and oncoprint^3^ to display mutation events in patient cohorts. Although these specialized visualizations offer deeper insights into data from specific domains, they suffer from limited customization and generalizability. In light of these challenges, there is growing demand for a generalized visualization paradigm that seamlessly integrates multiple types of plots, enabling the visualization of interrelations between features while maintaining an intuitive and unified interface.

To this end, we propose a visualization paradigm that introduces a novel and intuitive way to visualize complex datasets – the cross-layout (**Figure 1a**). At its core is a central main plot that focuses on representing the key feature of a dataset. Surrounding this central plot are secondary plots, incrementally added to each side. This arrangement forms a cross-like structure, hence the name “cross-layout visualization” (**Figure 1**). Each plot, central or secondary, allows the layering of additional features such as text or symbols. These layers add annotations and context, enhancing the understanding of the plots. For datasets with categorical axis, the paradigm allows incorporation of data-driven structure, for example, through hierarchical clustering showcasing similarities within and between data groups, adding a deeper analytical dimension (**Figure 1b-c**). Additionally, the paradigm offers versatility through concatenation and recursion: secondary plots can transform into central plots of new cross-layouts that are connected to the initial one, allowing for intricate and detailed visual representations of the data (**Figure 1d**). This approach makes complex data more accessible and interpretable and allows the visualization of cross-feature relationships.

**Figure 1:**
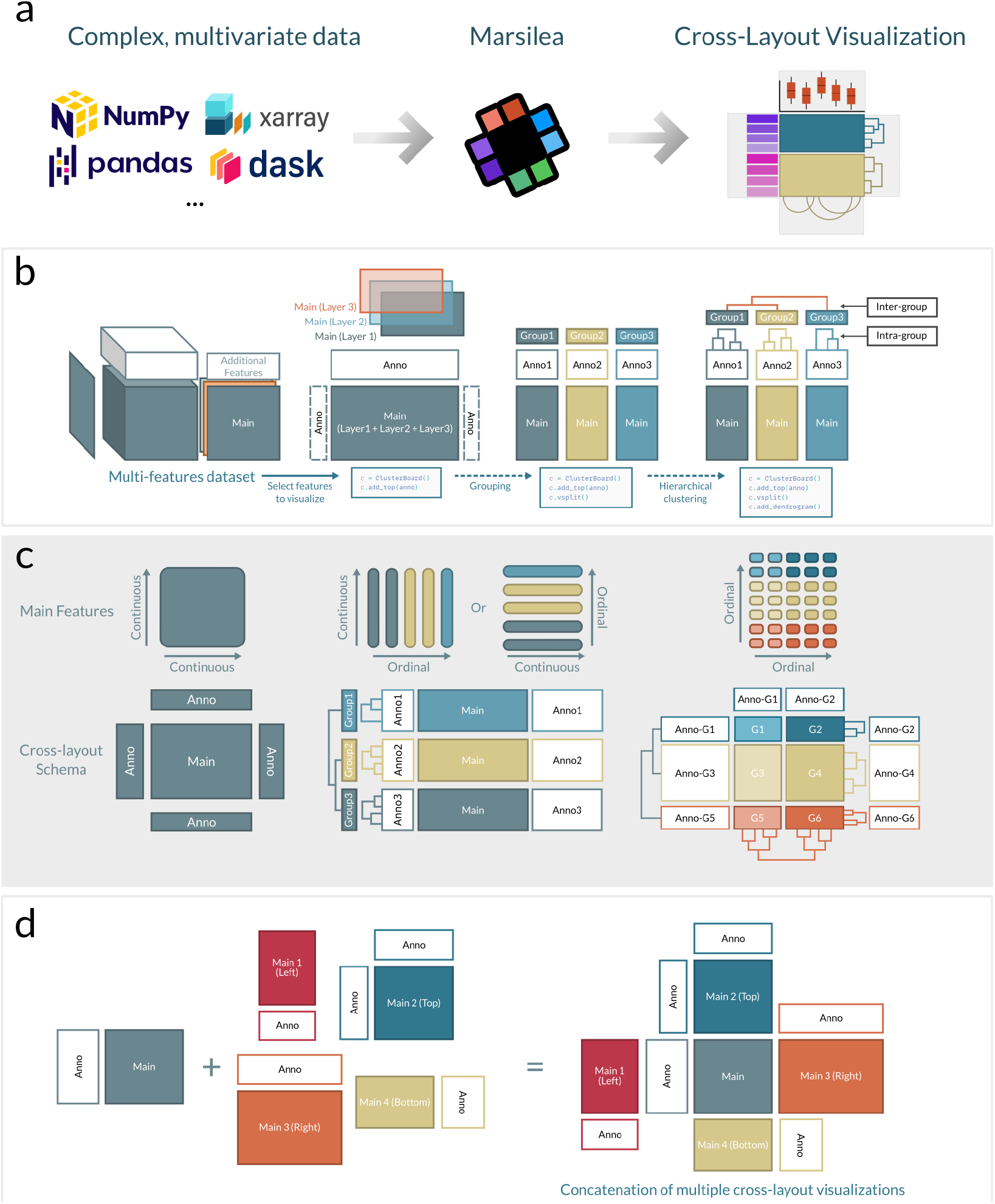
The novel concept of cross-layout visualization implemented in Marsilea. **a)** Conceptual illustration of the cross-layout visualization concept with a central plot and linked plots in its sides forming a cross-like shape. **b)** The assembly process of cross-layout visualization. **c)** Variants of cross-layouts adopted to different input data. **d)** Illustration of the versatility provided by concatenation of multiple cross-layouts.

To facilitate the creation and manipulation of cross-layout visualization, we built Marsilea, an easy-to-use Python library designed to simplify the creation of complex visualizations. Marsilea is built with modularity in mind, allowing users to add plot components incrementally as needed. Marsilea offers a diverse range of builtin plot types: four variants of heatmap, statistical plots like line plot, bar plot, violin plot, arc diagrams, text labels, and sequence logos (**Extended Figure 1**). One of Marsilea’s key strengths is its capacity for customization; users can easily implement and integrate their own plot types into the framework. Marsilea boasts compatibility with multiple input formats, from the primitive Python list to NumPy arrays and Pandas DataFrames, ensuring seamless integration into the data analysis pipelines (**Figure 1a**).

To demonstrate the broad applicability of the cross-layout paradigm and the intuitive usage of Marsilea, we create and extend existing complex figures in the domain of Biology. First, we leverage a widely used singlecell RNA expression dataset (PBMC3K) to create a complex heatmap visualizing marker genes of different cell types stratified by cell lineage (**Figure 2a**). We illustrate how, in a very intuitive manner, the user can assemble the figure with twelve lines of code and easily adjust its features, such as the plot size, the space between plots, the order, or the placing of different plots (**Extended Figure 2a**). These features illustrate Marsilea’s flexibility in layout arrangements and the simplicity of applying custom styles. Second, we use a large single-cell multi-omics dataset from COVID-19 patients containing proteomics and transcriptomics^4^ information to create a side-by-side heatmap comprising two cross-layouts concatenated. This plot enables visual comparison of two omics profiles (**Figure 2b**) and includes additional information such as cell meta clusters, cell abundance, and gene expression to enrich the context of the central features displayed. This type of plot cannot be generated with other high-level visualization frameworks in Python, such as seaborn or plotnine. Next, we demonstrate the recreation of specialized types of plots that often need dedicated tools for visualization: sequence motifs from the ggmsa tool^5^ (**Figure 2c**) and; set intersections from the IMDB top 1000 movie dataset, which are typically generated using the Upset tool (**Figure 2d**); additionally, we present the oncoprint plot, usually made with cBioPortal, jointly visualizing mutational and expression profiles (**Extended Figure 3a**), as well as a network diagram showing the characters’ relationships in ‘Les Misérables’ (**Extended Figure 3b**). Due to the generalizable approach of the cross-layout paradigm, Marsilea can generate these plots in an intuitive and unified manner.

**Figure 2:**
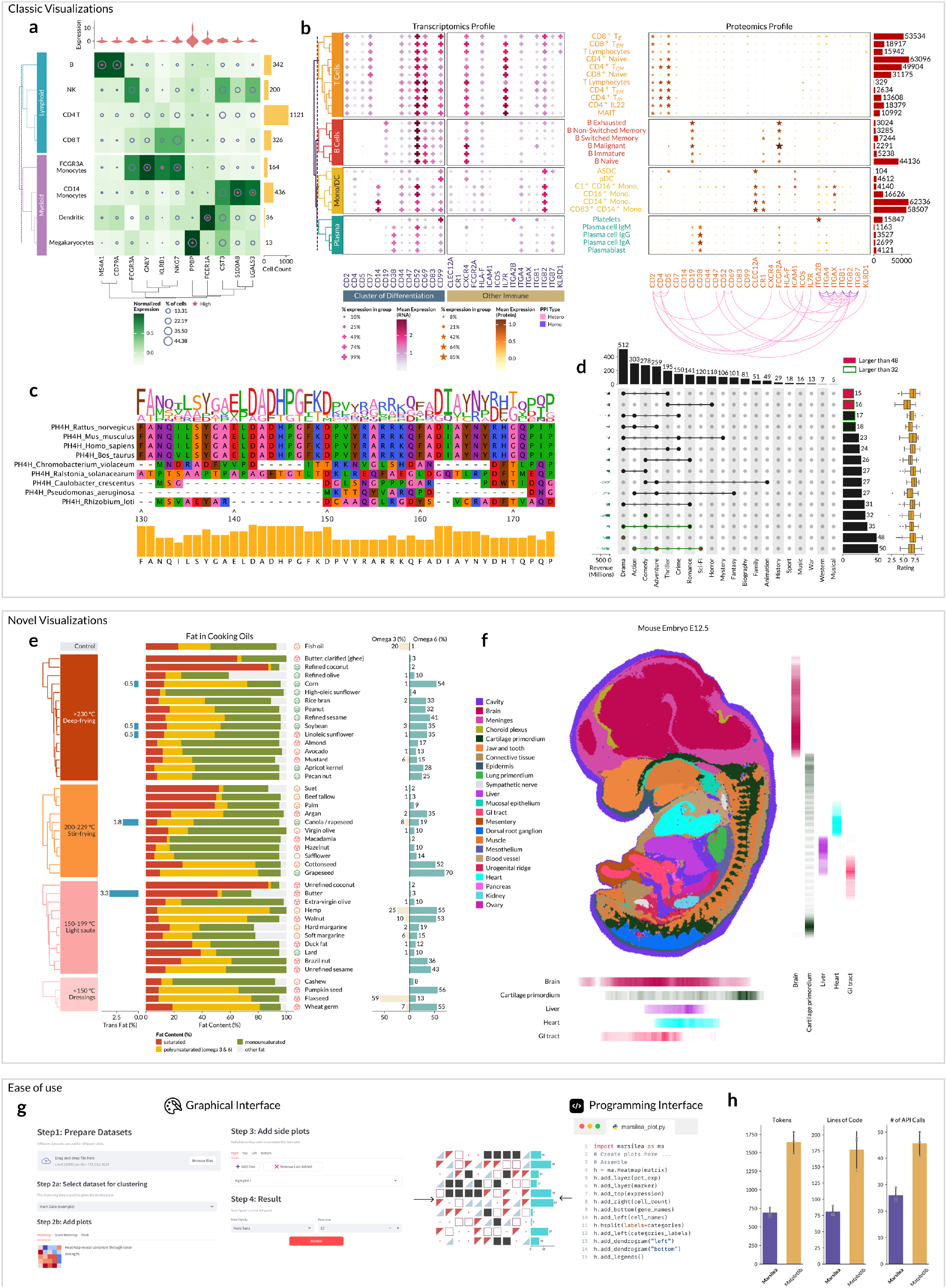
Demonstration of Marsilea’s capabilities reproducing existing figures and generating novel visualization types with ease. **a)** Reproduction of a common type of plot of cell-type aggregated gene expression from single-cell RNA sequencing. Note the use of the cross-layout and extensive use of annotations to the central plot. **b)** Visualization of multi-omics single-cell with two concatenated cross-layouts to enhance visual comparison of differences. **c)** Demonstration of a multiple sequence alignment plot for phenylalanine hydroxylase protein (PH4H) from nine species. **d)** Reproduction of a Upset plot showing genre overlap between Top 1000 IMDB movies. **e)** Novel visualization with multi-faceted information of daily cooking oils. **f)** Novel visualization displaying a spatial cell map of a mouse embryo with visual aids to discern cell types that encode in similar colors. **g)** Demonstration of the equivalence of graphic and programming interfaces of Marsilea. **h)** Benchmark of Marsilea against matplotlib showing considerable conciseness and simplicity.

Furthermore, to showcase the ability of the cross-layout paradigm to create novel visualizations that highlight the relationship between variables, we showcase novel visualizations for a variety of compositional data. In the first example, we utilize a dataset of fat content in 42 cooking oils (**Figure 2e**). The main feature, fat content, is depicted as stacked bars. We extend the stacked bar with additional features added incrementally as bar plots to the main feature, including the type of oil, symbols representing the subjective flavors, and the fraction of different healthy and unhealthy components in the oil: Omega 3 (Healthy), Omega 6 (Unhealthy), and transfat (Unhealthy). The plot was then grouped by the smoke point, showing the suitability of each oil for different cooking methods. Hierarchical clustering was performed to highlight fat content comparison, bringing oils with similar content closer together. Next, we create an enhanced version of the spatial cell map of a whole mouse embryo^6^. When visualizing the spatial distribution of cells in spatial omics data, it can be challenging to identify the location of specific cells in tissue when colors utilized in the legend are very similar. This is exacerbated for color-blind individuals (simulated color-blind image in **Extended Figure 4**). To address this issue, we extend the cell map by showing the density of specific cell types along the vertical and horizontal axes of the main plot (**Figure 2f**). The added density plots on the side assist in locating specific cell types within the embryo.

Marsilea provides multiple interfaces for users, catering to different user preferences and skill levels. It can be utilized programmatically through an object-oriented API for those comfortable with coding (**Figure 2g, Extended Figure 2a**) or through a no-code, web-based interface, making it accessible to a wider audience beyond just programmers (**Figure 2g, Extended Figure 2b**). The current web-based interface (http://marsilea.rendeiro.group) offers extensive customization, albeit not to the level conferred by the infinite customization of the programming interface. To assess the conciseness and simplicity of Marsilea’s code, we evaluated the implementation of the same visualization (**Figure 2e**) produced using Marsilea or its low-level counterpart, matplotlib (**Supplementary Methods**). We observed that Marsilea only demands about half the coding effort to produce identical visualizations (**Figure 2h**), suggesting it is significantly more user-friendly. Moreover, Marsilea’s layout adjustments are notably precise, requiring only a single line of code to modify plot location, size, and padding (**Extended Figure 2c**), which greatly facilitates going from individual plots to publication-ready figures for researchers.

In summary, the cross-layout paradigm is a simple way to create rich visualization with flexibility in composition and customization to unlock infinite possibilities of complex data visualization across scientific domains. With the flexibility and extensibility of Marsilea, we anticipate the integration of the cross-layout in various data analysis pipelines and research contexts, enabling scientists to create tailored visualizations that highlight cross-feature interactions with ease and in an intuitive manner.

## Data and Code Availability

Marsilea is open source and available on GitHub at https://github.com/Marsilea-viz/marsilea. A graphical user interface is available from http://marsilea.rendeiro.group. All datasets used in this work are publicly available at https://github.com/Marsilea-viz/marsilea-data, with the data sources documented in the repository. The datasets and the code to recreate the examples in the figure can be found in the GitHub repository and the documentation website at https://marsilea.readthedocs.io.

### Contributions

YZ proposed the idea of a cross-layout and developed a demo. YZ and ZZ developed the Python package and built the documentation website together. YZ developed the web application for Marsilea. YZ and EC drafted the manuscript together. AFR edited the manuscript and figures. AFR and EC supervised the research. All authors read and approved the final manuscript.

## Acknowledgments

We thank the open-source community efforts of building matplotlib^7^, seaborn^8^, numpy^9^, and pandas^10^, which made this project possible. This work was supported by the University of Macau [MYRG2020-00100-FHS, MYRG2022-00204-FHS, MYRG-GRG2023-00189-FHS-UMDF]; and Macau Science and Technology Development Fund [0011/2019/AKP, 0137/2020/A3]. YZ and AFR are supported by Angelini Ventures S.p.A. Rome, Italy.

## Supplementary Methods

### Benchmarking procedure

We asked three individuals to implement the cooking oils example in Figure 2e. Their code files were first formatted using ruff (https://github.com/astral-sh/ruff) with default settings and then measured with the number of tokens, lines of codes (no blank lines, no comment lines), and the number of API calls. The benchmark scripts can be found in the GitHub repository of Marsilea (https://github.com/Marsileaviz/marsilea).

## Extended Data Figures

**Extended Data Figure 1:**
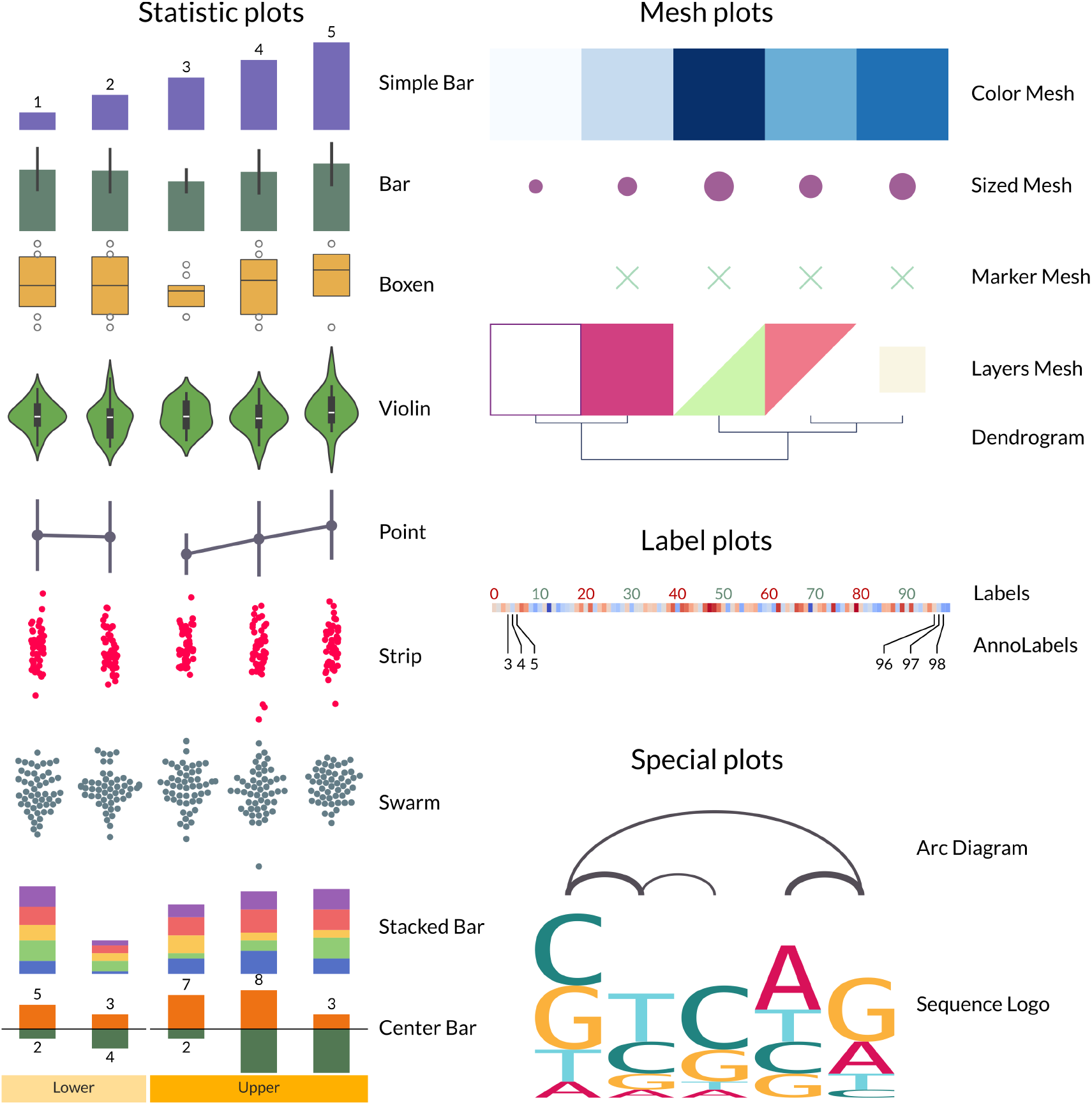
Demonstration of the basic building blocks of Marsilea.

**Extended Data Figure 2:**
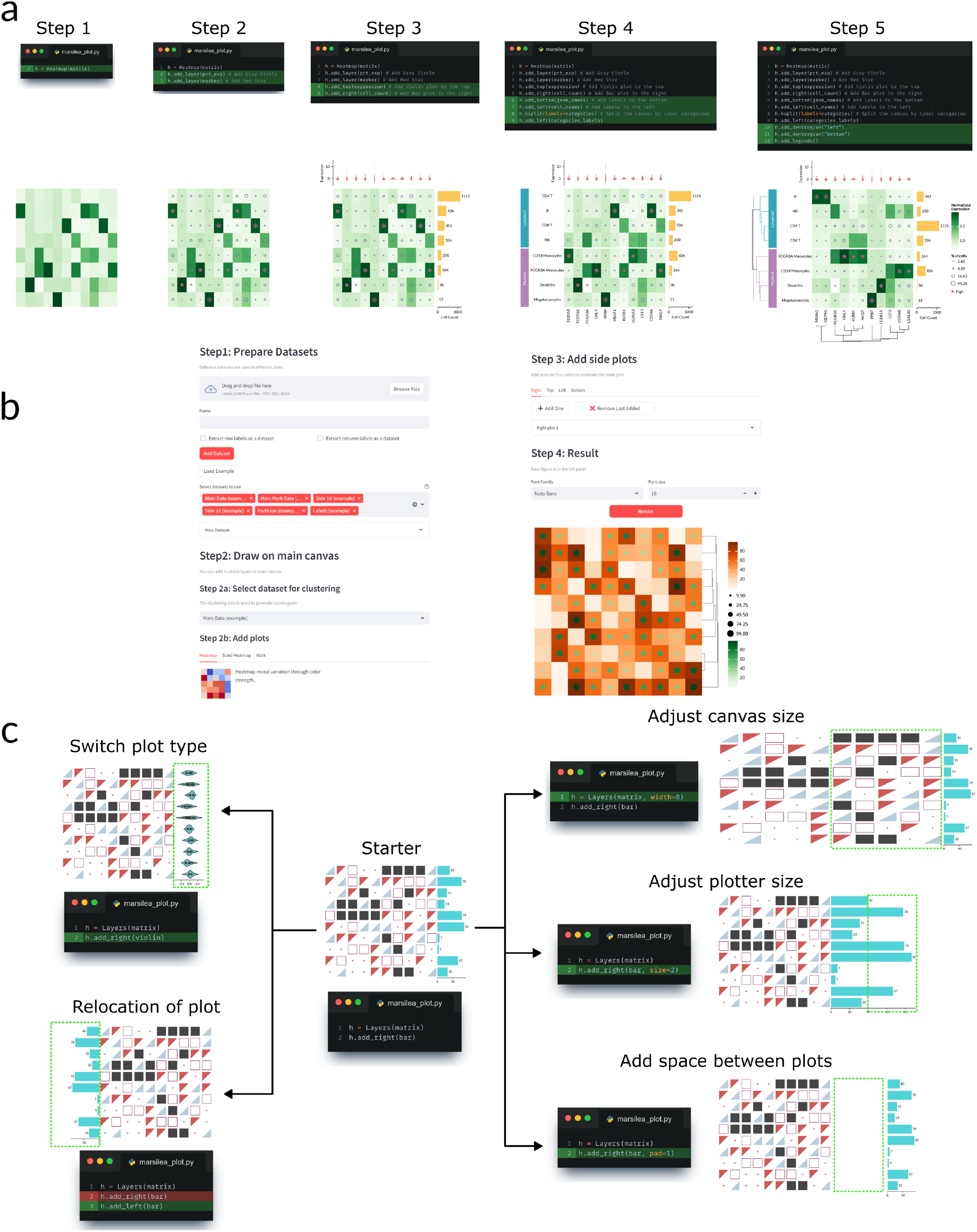
Demonstration of the **a)** programmatic and **b)** web-based interfaces of Marsilea. **c)** Demonstration of the simplicity to make layout adjustment in Marsilea.

**Extended Data Figure 3:**
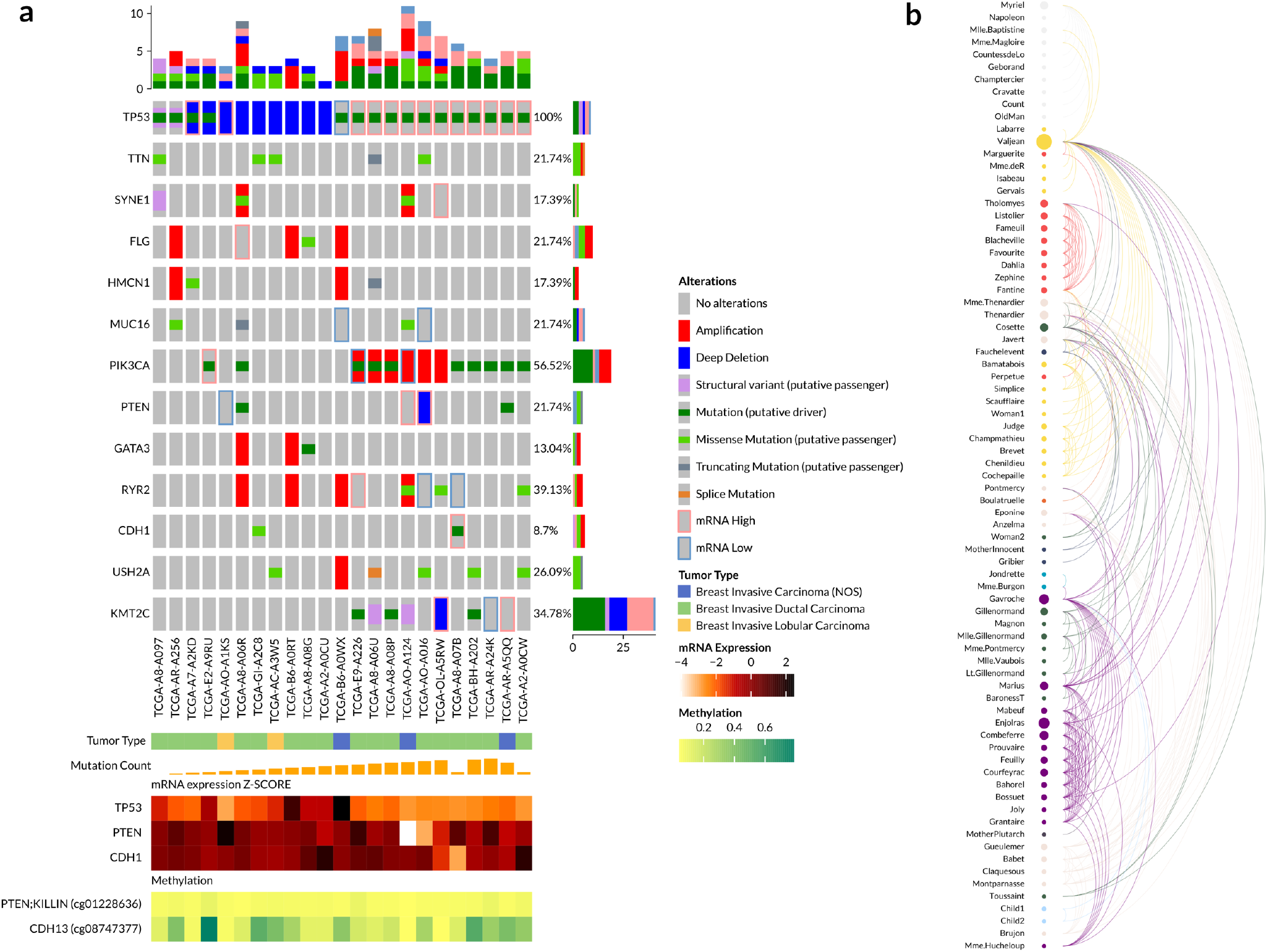
Demonstration of different visualizations created by Marsilea. **a)** Oncoprint plot visualize a subset of TCGA breast cancer mutation, mimicking the original style from cBioPortal. **b)** Arc diagram showing the character relationships in Les Misérables.

**Extended Data Figure 4:**
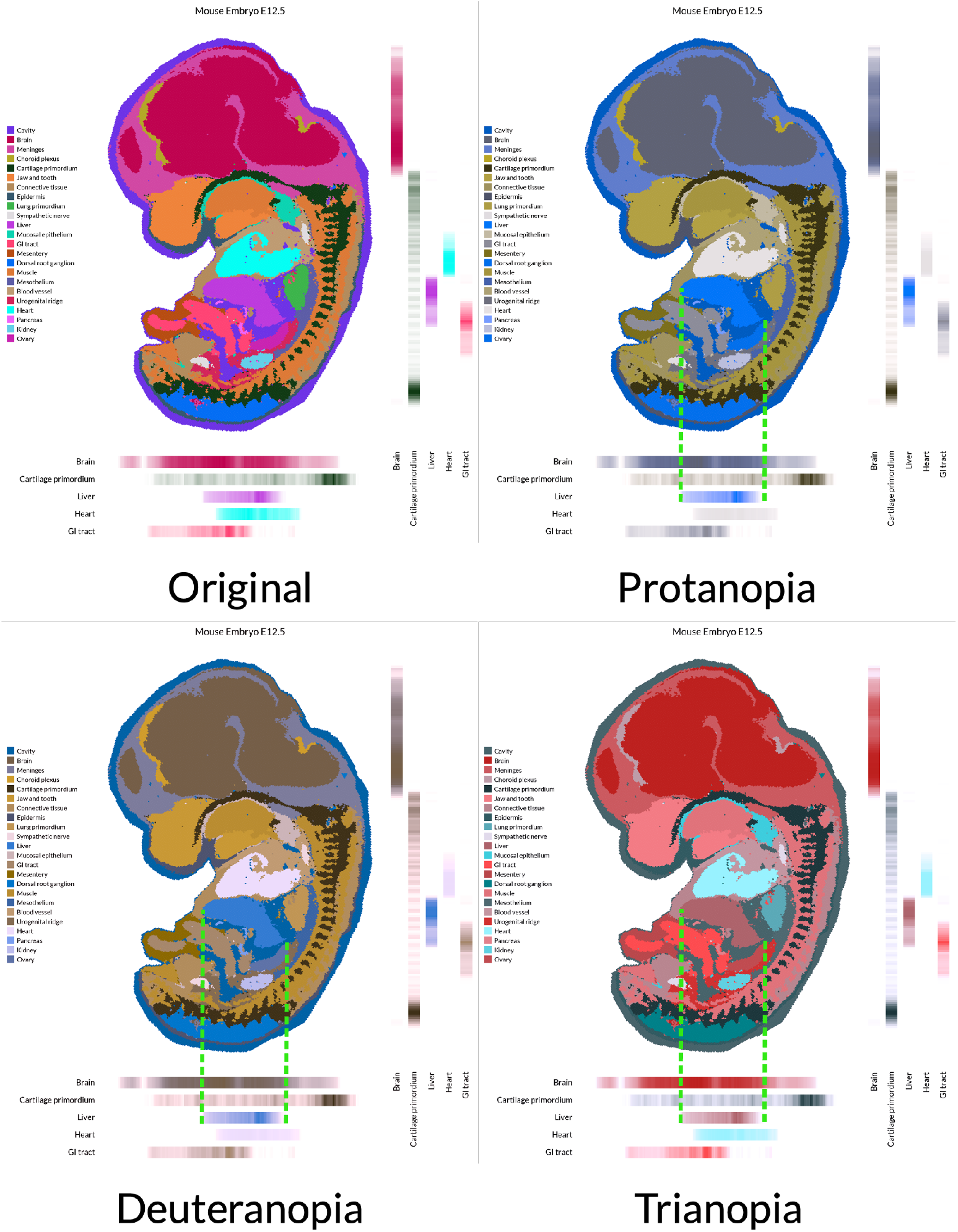
Simulation of three variants of color blindness.

## References

1. McInnes, L., Healy, J. & Melville, J. UMAP: Uniform Manifold Approximation and Projection for Dimension Reduction. arXiv [stat.ML] (2018).

2. Gu, Z., Eils, R. & Schlesner, M. Complex heatmaps reveal patterns and correlations in multidimensional genomic data. Bioinformatics 32, 2847–2849 (2016).

3. Gao, J. et al. Integrative analysis of complex cancer genomics and clinical profiles using the cBioPortal. Sci. Signal. 6, 1 (2013).

4. Stephenson, E. et al. Single-cell multi-omics analysis of the immune response in COVID-19. Nat. Med. 27, 904–916 (2021).

5. Zhou, L. et al. ggmsa: a visual exploration tool for multiple sequence alignment and associated data. Brief. Bioinform. 23, (2022).

6. Chen, A. et al. Spatiotemporal transcriptomic atlas of mouse organogenesis using DNA nanoball-patterned arrays. Cell 185, 1777–1792.e21 (2022).

7. Hunter. Matplotlib: A 2D Graphics Environment. Comput. Sci. Eng. 9, 90–95 (2007).

8. Waskom, M. seaborn: statistical data visualization. J. Open Source Softw. 6, 3021 (2021).

9. Harris, C. R. et al. Array programming with NumPy. Nature 585, 357–362 (2020).

10. Mc Kinney, W. Pandas: A foundational python library for data analysis and statistics. https://www.dlr.de/sc/portaldata/15/resources/dokumente/pyhpc2011/submissions/pyhpc2011_submission_9.pdf (2011).

